# Overlap and divergence of neural circuits mediating distinct behavioral responses to sugar

**DOI:** 10.1101/2023.10.01.560401

**Authors:** Ruby V. Jacobs, Crystal X. Wang, Fiorella V. Lozada-Perdomo, Lam Nguyen, Julia U. Deere, Hannah A. Uttley, Anita V. Devineni

## Abstract

A single sensory cue can elicit diverse behavioral responses. For example, the taste of sugar robustly promotes feeding^1, 2^ but also influences other behaviors, such as altering locomotor patterns to maximize food-finding^3, 4^ or conferring a rewarding value onto associated contexts or cues.^5–7^ Here, we investigate how sweet taste elicits multiple appetitive behaviors in *Drosophila*. Are different sugar-evoked behaviors coordinately regulated? At what point does the sugar circuit diverge into different pathways that drive distinct behaviors? We first established an optogenetic paradigm to study the effects of sugar taste on locomotion, spatial preference, and associative learning. We then tested how different sugar-evoked behaviors were modulated by internal and external factors, including hunger, diet, or the presence of an aversive taste. Different behaviors were generally modulated in similar ways, but we also observed some differences that reveal selective modulation of specific behavioral pathways. Finally, we investigated where the sugar taste circuit diverges into different behavioral pathways. A recent study identified a sensory-motor circuit comprising five layers of neurons that drives the initiation of feeding in response to sugar.^8^ By individually manipulating each of these neurons, we show that circuits mediating different innate responses to sugar are partially overlapping and begin to diverge at the level of second- and third-order neurons, whereas circuits for innate versus learned behaviors may diverge at the first synapse. Connectomic analyses reveal distinct subcircuits that mediate different behaviors. Together, these studies provide insight into how neural circuits are organized to elicit diverse behavioral responses to a single stimulus.

## RESULTS

### Activating sugar-sensing neurons elicits locomotor suppression, positional preference, and appetitive learning

Studies of sugar taste in *Drosophila* have primarily focused on measuring feeding initiation or food consumption.^8–17^ However, sugar also influences other behaviors related to feeding. The taste of sugar suppresses locomotion to enable flies to feed,^18, 19^ causes flies to preferentially reside in sugar-containing areas,^19, 20^ and drives appetitive learning for odors associated with sugar.^21, 22^ We established an optogenetic paradigm to study these behaviors. We used the light-gated channel Chrimson^23^ to optogenetically activate sugar-sensing neurons across the body using *Gr64f-Gal4*.^10^ These include neurons in the legs and the labellum, the distal segment of the proboscis (the fly’s feeding organ). The optogenetic approach allows us to precisely control the timing and duration of taste stimulation, as well as avoiding potential confounding effects resulting from sugar consumption during the assay (e.g., satiation). Because feeding responses to sugar are enhanced by food deprivation,^11, 24, 25^ we compared fed and starved flies in each assay.

We first tested how the activation of sugar-sensing neurons affects locomotion. We tracked flies’ movement while delivering 5 sec red light stimulation at three different intensities (“low”, “medium”, and “high”), which corresponded to 4, 20, and 35 µW/mm^2^ (Figure 1A). Activating sugar-sensing neurons strongly suppressed locomotion at all light intensities (Figure 1B-C), consistent with previous studies using natural sugar.^18, 19^ Both forward velocity and turning were suppressed at light onset, and the fraction of flies moving decreased to nearly zero (Figure 1B-C and S1A). In fed flies, this locomotor suppression was transient and flies resumed movement after 2-3 seconds even though the light stimulation remained on, perhaps because flies quickly determine that there is no food source actually present. The locomotor suppression was strongly enhanced and prolonged in flies that were starved for one day, although their speed also began to recover toward the end of the light stimulus (Figure 1C).

**Figure 1.**
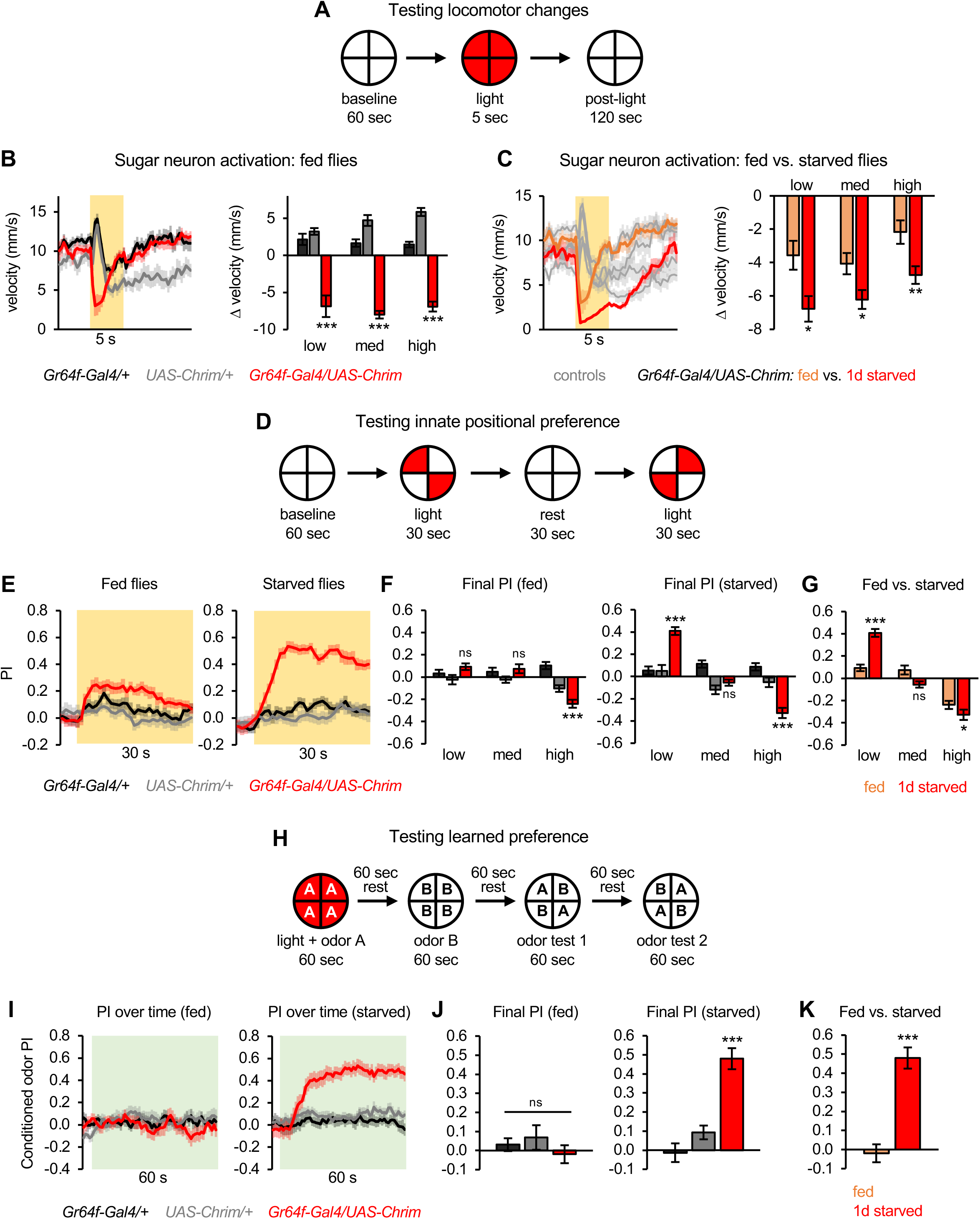
Activating sugar-sensing neurons elicits locomotor suppression, positional preference, and appetitive learning. (A) Schematic of protocol to test the effect of sugar neuron activation on locomotion. (B-C) Activating sugar-sensing neurons elicited locomotor suppression that was enhanced by starvation (n = 11-12 trials). 5 sec light stimulation was used at 3 different intensities (low, medium, high). Left graphs show forward velocity over time for low light intensity. In all figures, yellow shading indicates light on. Bar graphs on the right quantify the light-evoked change in forward velocity. (B) Comparison of experimental and control genotypes in the fed state. (C) Comparison of fed and one-day starved flies. Bar graphs show change in velocity during the first 1 sec of light onset (B) or the entire light period (C); changes during the first 1 sec were similar in fed and starved flies. (D) Schematic of protocol to test the effect of sugar neuron activation on innate positional preference (adapted from Aso et al., 2014). Light preference was quantified by the preference index (PI): (# flies in light quadrants – # flies in non-light quadrants) / total # flies. (E-G) Activating sugar-sensing neurons elicited significant positional preference in starved flies, but not fed flies (n = 22-24 trials, 11-12 sets of flies). (E) PI over time for low intensity light presentation. (F-G) Final PI for control versus experimental flies (F) or fed versus one-day starved experimental flies (G). For all figures, final PI represents the average PI over the last 5 sec of light or odor presentation. (H) Schematic of protocol to test the effect of sugar neuron activation on learned odor preference (adapted from Aso & Rubin, 2016). Learned preference was quantified as the PI for the conditioned odor during the test periods: (# flies in CS+ quadrants – # flies in CS-quadrants) / total # flies. Data from test 1 and 2 are combined. (I-K) Activating sugar-sensing neurons elicited learned preference in starved flies, but not fed flies (n = 16-24 trials, 8-12 sets of flies). (I) PI during the odor test period. In all figures, green shading indicates odor on. (J-K) Final PI (average over last 5 sec of odor test) for control versus experimental flies (J) or fed versus one-day starved experimental flies (K). For all figures: *p<0.05, **p<0.01, ***p<0.001, ns = not significant (p>0.05). Unless otherwise specified, two groups were compared using unpaired t-tests and more than two groups were analyzed using one-way ANOVA followed by Dunnett’s test comparing experimental flies to each control.

We next tested whether flies prefer residing in areas where they receive sugar neuron activation, similar to their positional preference for natural sugar.^19, 20^ We presented light in two opposing quadrants of the arena (Figure 1D) and quantified the flies’ preference for residing in the light quadrants. Activating sugar neurons at low light intensity elicited strong positional preference in one-day starved flies, but only a weak, non-significant effect in fed flies (Figure 1E-G). Unexpectedly, the attraction diminished at medium light intensity and shifted to aversion at high intensity (Figure 1F-G). This high intensity aversion has been previously observed^26^ and may reflect opposing functions of different *Gr64f-Gal4*-expressing cells or a compensatory change in the sugar circuit (see Discussion). Although locomotor suppression (Figure 1B-C) could lead to positional preference (Figure 1E-G) by making flies more likely to remain in a certain area, these two behaviors are clearly dissociable. Locomotor suppression was observed at all light intensities in both fed and starved flies (Figure 1B-C), whereas positional preference was only observed in starved flies at the lowest intensity (Figure 1E-G).

In addition to its effects on innate behavior, sugar can be used as a reinforcement cue for associative learning.^21, 22^ We tested whether optogenetic activation of sugar-sensing neurons could drive learned preference toward an odor. Our learning assay follows previously established protocols^27^: we delivered one odor (the conditioned stimulus, CS+) while activating sugar neurons, then presented a different odor (the CS-) without neuronal activation, and finally allowed the flies to choose between the CS+ and CS-odors presented in opposing quadrants (Figure 1H). We observed learned preference for the CS+ in one-day starved flies, but not fed flies (Figure 1I-K). Neuronal activation in these experiments was initially performed using low light intensity because only low light elicited innate preference (Figure 1E-G), but stimulation with a higher intensity (30 µW/mm^2^) also caused learned preference in starved flies (Figure S1B).

These results show that the activation of sugar-sensing neurons drives behaviors evoked by natural sugar: locomotor suppression, positional preference (at low light intensity), and appetitive learning. Because the locomotion and positional preference assays were conducted with continuous light stimulation, we also tested 50 Hz pulsed light stimulation and observed the same behavioral effects, including attraction only at low light intensity (Figure S1C-D). In addition, we tested a second Gal4 line labeling sugar-sensing neurons, *Gr5a-Gal4.*^10^ Similar to activation with *Gr64f-Gal4*, activation of sugar neurons using *Gr5a-Gal4* in starved flies caused strong locomotor suppression at all light intensities and positional preference at only the lowest intensity (Figure S1E-F). We did not observe learned preference using *Gr5a-Gal4* (Figure S1G), which could be related to the fact that *Gr64f-Gal4* labels more sugar neurons than *Gr5a-Gal4*.^10^ It is possible that stronger learning protocols (e.g., with repeated pairings) could elicit learning using this line.

### Hunger selectively modulates specific taste pathways and behavioral responses

Our results show that all of the behavioral responses to sugar neuron activation were enhanced in starved flies, but to varying degrees: fed flies did not show any associative learning (Figure 1I-K), but they showed a weak positional preference (Figure 1E) and strong locomotor suppression (Figure 1B-C). Thus, hunger state strictly gates certain sugar-evoked behaviors but exerts a graded effect on other behaviors, suggesting that hunger acts downstream of sensory neurons to differentially modulate each behavioral pathway.

The behavioral effects of activating sugar-sensing neurons represent exactly the opposite effects from those observed when bitter-sensing neurons are activated,^28^ indicating that appetitive and aversive tastes regulate the same behaviors in opposing directions. We wondered how hunger modulates the same behaviors in response to opposing taste modalities. We considered three models: 1) Hunger potentiates responses to all tastants, thus enhancing both appetitive responses to sugar and aversive responses to bitter. 2) Hunger biases behavior toward appetitive responses, thus enhancing appetitive responses to sugar while suppressing aversive responses to bitter. Studies of feeding behavior support this model.^13, 29^ 3) Hunger selectively modulates behavioral responses to sugar but not bitter taste, reflecting the idea that hungry animals should consume more calories while retaining the ability to reject contaminated food.

To distinguish between these models, we compared the effects of bitter neuron activation in fed and starved flies. In contrast to the strong effect of hunger on sugar-evoked behaviors, hunger had a much weaker effect on behaviors evoked by bitter neurons. Hunger modestly enhanced aversive learning with bitter but did not affect bitter-evoked locomotor changes or positional aversion, even when flies were starved for two days (Figure S1H-I). These results show that hunger acts selectively to modulate specific taste pathways and behavioral responses.

### Aversive taste overrides all sugar-evoked behaviors

We next asked how sugar-evoked behaviors are affected by simultaneously activating bitter neurons, which drive opposing responses. The opposing effects of sugar and bitter activation could be linearly integrated and therefore result in an intermediate response, or the effects of one pathway could dominate over the other. We also wondered whether sugar and bitter inputs are integrated in similar or different ways to regulate distinct behaviors.

In the locomotor assay, co-activating sugar and bitter neurons elicited effects similar to activating bitter neurons alone, causing a strong increase in forward velocity and turning (Figure 2A-B). Thus, the locomotor enhancement driven by bitter neurons overrides the locomotor suppression driven by sugar neurons. However, closer examination of the light onset period revealed that sugar and bitter neuron co-activation caused very brief locomotor suppression, resulting in a longer delay to reach peak velocity and turning as compared to activation of bitter neurons alone (Figure 2A-B). This trend was not significant for peak velocity (Figure 2A; p = 0.069 for starved flies), but was robust and highly significant for peak turning (Figure 2B). The delay to peak turning was also enhanced in starved flies compared to fed flies (Figure 2B).

**Figure 2.**
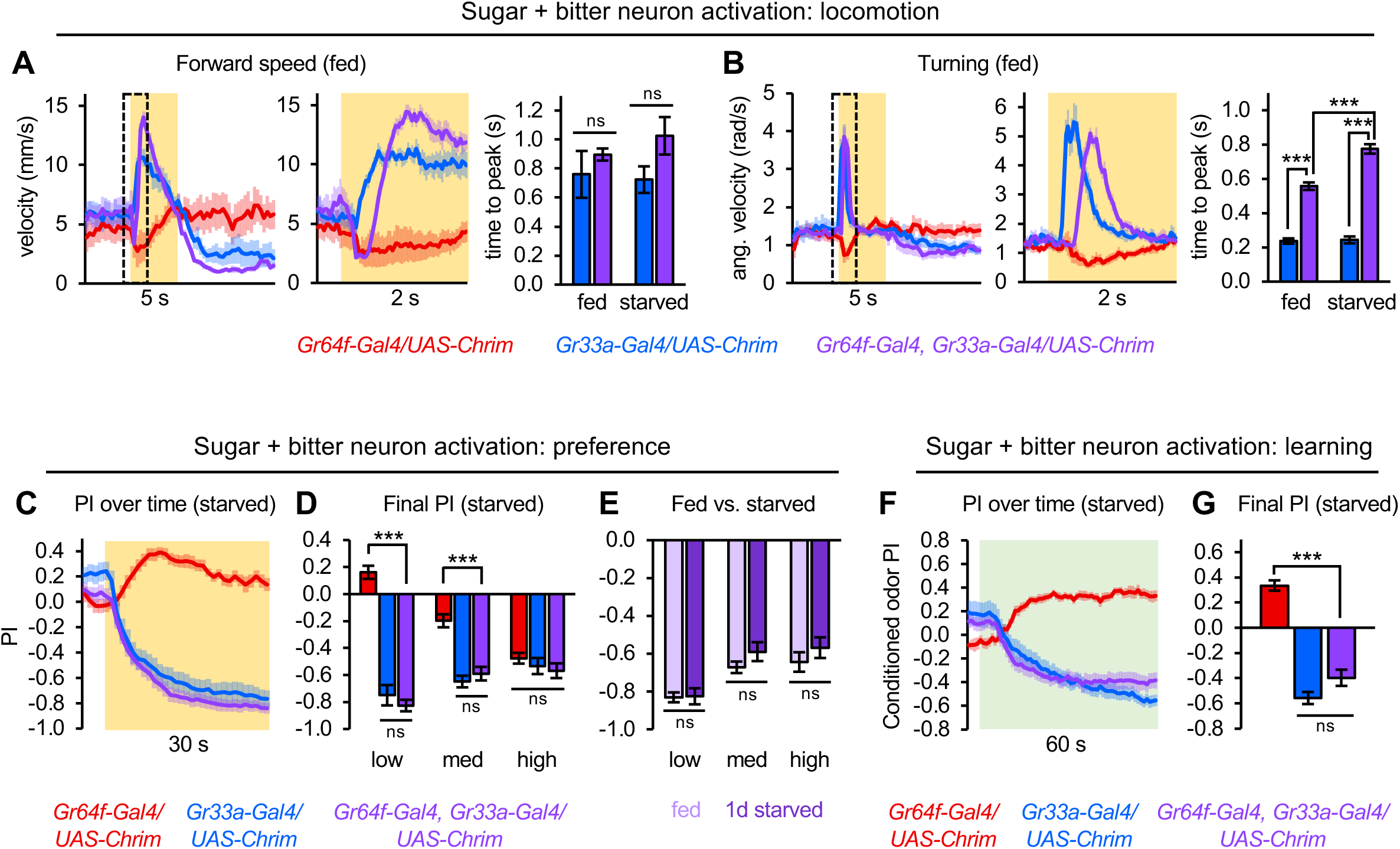
All behaviors elicited by sugar neurons are potently suppressed by co-activation of bitter neurons. (A-B) Co-activating sugar- and bitter-sensing neurons caused brief locomotor suppression followed by a strong increase in forward speed and turning (n = 9-10 trials). Traces show behavior of fed flies with low intensity light stimulation. Right traces show zoomed in behavior around light onset (boxes shown in left graphs). Bar graphs quantify the time to peak velocity following light onset for fed versus one-day starved flies when bitter neurons were activated alone (blue) or along with sugar neurons (purple). (C-E) Co-activating sugar- and bitter-sensing neurons elicited strong positional aversion in fed and one-day starved flies (n = 18-20 trials, 9-10 sets of flies). Effects with low light intensity are shown in panel C. (F-G) Co-activating sugar- and bitter-sensing neurons elicited learned aversion in one-day starved flies (n = 20 trials, 10 sets of flies). *Gal4/+* and *UAS/+* controls are not shown in this figure but were tested alongside the experimentals and behaved similarly to the controls shown in Figures 1 and S1.

These results suggest that when sugar and bitter neurons are co-activated, the two inputs are integrated differently across time: sugar input drives locomotor behavior during the earliest time points (and hunger prolongs its influence), while bitter input dominates throughout the rest of the stimulation period. Under natural conditions, these effects may lead to sequential behaviors when flies encounter a mixture of sugar and bitter: the taste of sugar causes flies to briefly stop and consider feeding, but sufficiently strong bitter taste overrides this program and causes them to run away.

In the positional preference and learning assays, co-activation of sugar and bitter neurons elicited a similar degree of avoidance as activating bitter neurons alone (Figure 2C-G). Thus, the innate or learned aversion elicited by bitter taste completely dominates over the attraction elicited by sugar taste. These results show that across multiple behaviors, bitter input overrides sugar input to drive aversive behaviors.

### A high-sugar diet reduces behavioral responses to sugar neuron activation

The experiments described above show that multiple sugar-evoked behaviors are enhanced by hunger and suppressed by bitter taste, revealing the coordinated regulation of different behavioral responses. We next asked how these responses are affected by a high-sugar diet. Diet composition alters taste perception in many species, including humans, with dietary sugar being inversely correlated with sensitivity to sweet taste.^15, 17, 30–37^ Flies fed on a high-sugar diet show blunted responses of sugar-sensing neurons and reduced proboscis extension to sugar, a behavior that represents the initiation of feeding.^15, 17, 38, 39^ However, these flies increase their sugar consumption over several days and eventually eat far more than control flies, perhaps representing a compensatory response to their dulled sweet perception.^17^ We therefore wondered how a high-sugar diet influences other behavioral responses to sugar that occur on intermediate timescales. The behavioral responses we are measuring are not as immediate as proboscis extension but still represent short-term responses measured over seconds to minutes rather than days.

We compared flies raised on a control diet (standard fly food) or a high-sugar diet (fly food with 30% sucrose) for 7 days. We found that the high-sugar diet blunted behavioral responses to sugar neuron activation. Flies on a high-sugar diet showed more transient locomotor suppression, an effect that was apparent in both fed and starved flies (Figure 3A-B and S2A). Diet did not significantly affect baseline locomotion for any individual genotype, although in fed flies the high-sugar diet tended to decrease locomotion (Figure S2B). In both the innate and learned preference assays, flies on the control diet showed significant preference whereas flies on the high-sugar diet did not (Figure 3C-D). The deficit in learned preference is consistent with a recent study testing learning with natural sugar.^40^

**Figure 3.**
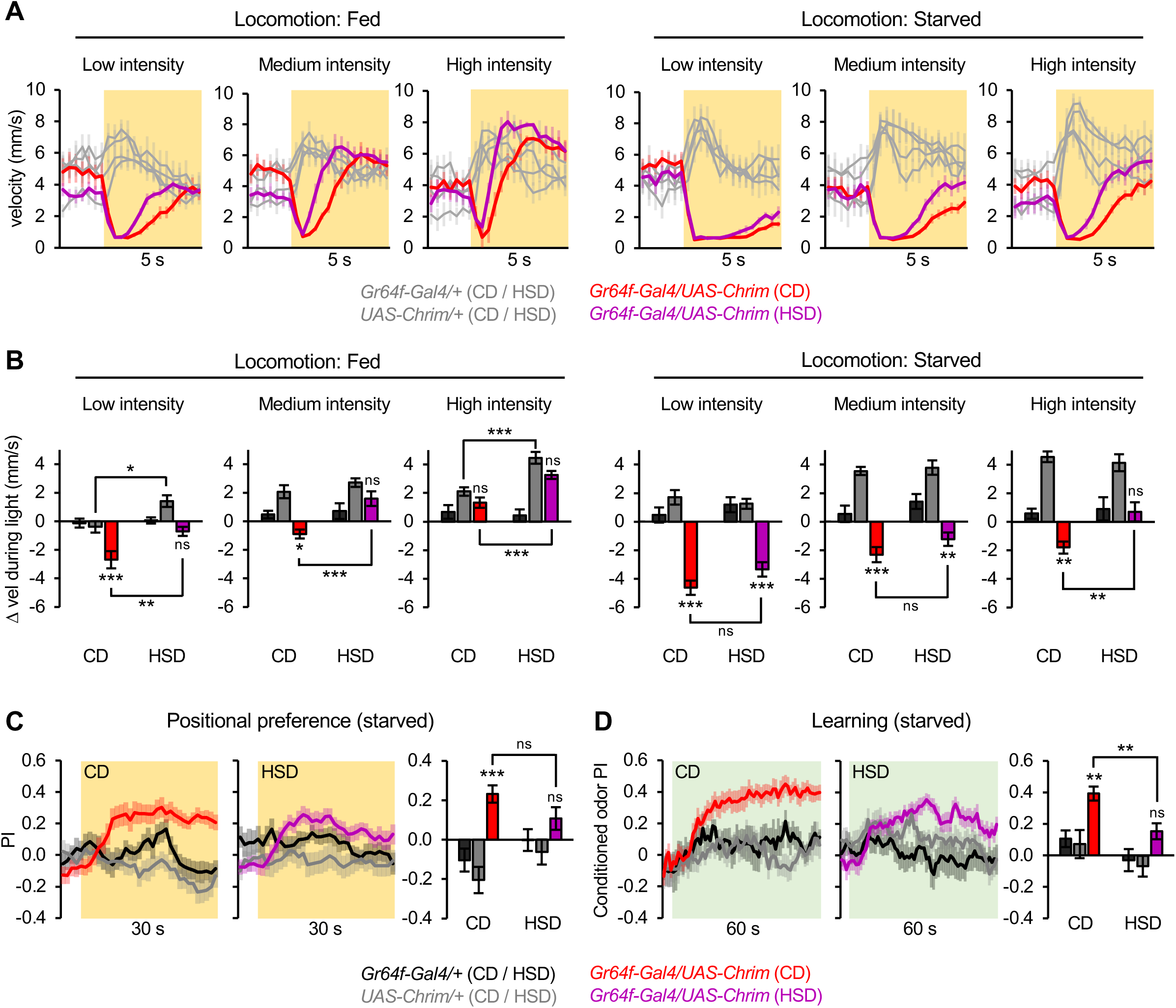
A high-sugar diet reduces behavioral responses to sugar neuron activation. (A) Locomotor suppression elicited by sugar neuron activation was more transient for flies raised on a high-sugar diet (HSD) compared to flies raised on a control diet (CD) (n = 7-10 trials). Flies were either fed (left) or one-day starved (right). The traces are zoomed in compared to other graphs (e.g., Figure 1) in order to show differences in how long the locomotor suppression lasts during the 5 sec light stimulation. (B) Change in velocity during the entire light period was quantified from the data in panel A. The change in velocity at light onset is shown in Figure S2A. (C-D) Sugar neuron activation at low light intensity elicited significant positional preference (C; n = 14-20 trials, 7-10 sets of flies) and learned preference (D; 10-18 trials, 5-9 sets of flies) in one-day starved flies raised on a CD, but not a HSD. All statistics in this figure represent the results of two-way ANOVA followed by Dunnett’s post-tests comparing experimental flies to each control or Bonferroni’s post-tests comparing CD and HSD.

Taken together with previous studies testing proboscis extension,^17, 39^ these results suggest that excess dietary sugar globally dampens short-term behavioral responses to sugar. Because we used optogenetic stimulation of sugar neurons, which bypasses sugar receptor activation, our results additionally demonstrate that a high-sugar diet decreases sugar sensitivity by acting downstream of the sugar receptors.

### Overlap and divergence of pathways for different sugar-evoked behaviors

We next investigated the neural circuits that underlie these diverse behavioral responses to sugar taste. At what point does the sugar circuit diverge into different pathways that drive different behaviors? This divergence could occur early in the circuit, potentially at the first synapse, or it could occur after multiple layers of sensory processing. Little is known about the neural circuits that mediate behavioral responses to sugar, with the exception of proboscis extension: a recent study by Shiu, Sterne et al. identified a core circuit within the subesophageal zone (SEZ) that drives proboscis extension in response to sugar.^8^ This circuit connects sugar-sensing neurons to proboscis motor neurons via three layers of interneurons, and each interneuron evokes proboscis extension when individually activated.^8^ We asked whether these interneurons also drive other sugar-evoked behaviors, which would reveal where the sugar circuit diverges into separate pathways for regulating different behaviors.

We tested the effect of individually activating each of the known interneurons in the proboscis extension circuit, which include 7 types of second-order neurons, 2 types of third-order neurons, and one fourth-order/premotor neuron (Figure 4A). We targeted each interneuron type using previously characterized, highly specific split-Gal4 lines,^8, 41^ and we verified the expression pattern of each line using immunostaining (not shown). Each neuronal “type” comprises a single neuron per hemisphere with the exception of Bract, which consists of 2 cells per hemisphere, but only one cell appears to be labeled in the split-Gal4 lines. We confirmed that optogenetic activation of each interneuron elicits proboscis extension, as previously demonstrated,^8^ for all but one line (Usnea) (Figure S3A). The lack of an effect with Usnea could result from differences in our experimental conditions (e.g., light intensity, immobilization method). We initially tested one split-Gal4 line per neuronal type; if phenotypes were borderline we then tested a second line (if available). Flies were one-day starved to maximize our chance of observing behavioral effects.

**Figure 4.**
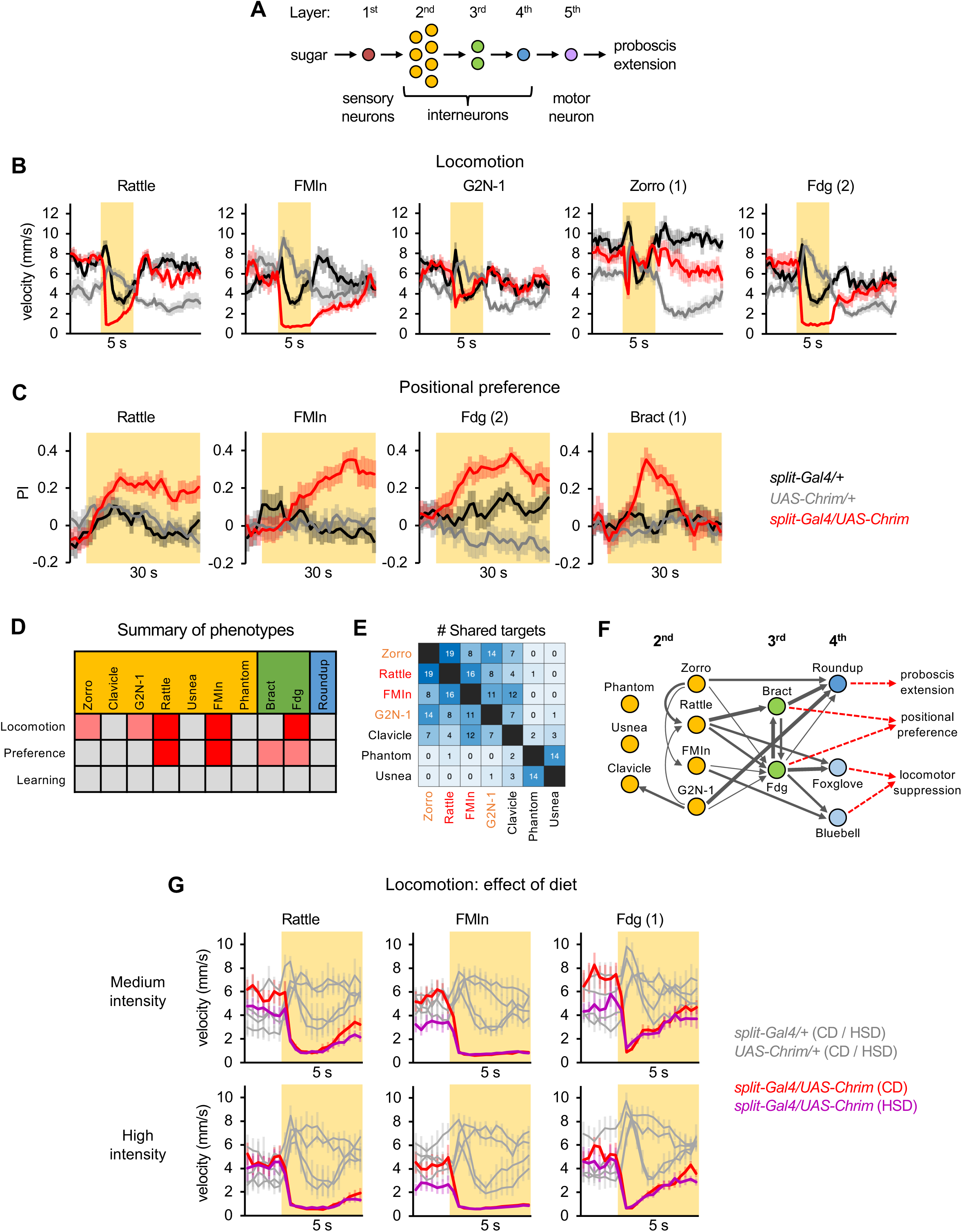
Overlap and divergence of pathways for different sugar-evoked behaviors. (A) Schematic of the known neural circuit that drives proboscis extension to sugar. Other neurons in the sugar pathway have been identified in the connectome and may contribute to this response (see Shiu, Sterne et al., 2022) but their roles have not been functionally validated due to the lack of Gal4 lines. (B) 5 of 10 interneuron types caused locomotor suppression when individually activated (n = 7-12 trials). Effects with medium light intensity are shown for all neurons except Zorro, for which high intensity light stimulation is shown. The locomotor suppression elicited by Zorro was extremely transient, but was stronger than it appears in this figure because these traces show binned velocity over 0.33 sec. Locomotor responses for all interneuron types at each light intensity are shown in Figure S3B. (C) 4 of 10 interneuron types caused positional preference when activated (n = 16-24 trials, 8-12 sets of flies), although the final preference was only significant for Rattle and FMIn. Effects with medium light intensity are shown for all neurons except Bract, which only showed an effect at high intensity. Preference effects for all interneuron types at each light intensity are shown in Figure S3C. (D) Summary of behavioral results with interneurons. Red shading indicates a significant and strong phenotype; light red shading indicates a phenotype that was relatively weak (Zorro and G2N-1 locomotion) or not statistically significant at the usual timepoints analyzed (Fdg and Bract preference). (E) Heatmap showing the number of shared targets for each pair of second-order neurons. Neuron names are colored red if they affected both locomotion and preference or orange if they only affected locomotion. (F) Schematic of sugar circuit showing connections between neurons, based on connectome analysis, and their relationship to behavior. Arrow thickness represents connection strength: thin = 5-9 synapses; medium = 10-19 synapses; thick = 20+ synapses. Two recently described fourth-order neurons, Foxglove and Bluebell, are included in the diagram but were not functionally studied in this paper. For simplicity, the diagram omits predicted inhibitory connections (from Phantom and Usnea), feedback connections from 3^rd^ to 2^nd^ order neurons, and weak connections from FMIn to Foxglove and Rattle to Bluebell that are only observed in one hemisphere. (G) Locomotor suppression elicited by activating second- or third-order interneurons did not differ between flies raised on a control diet (CD) or high-sugar diet (HSD) (n = 6-11 trials). Effects with medium (top row) or high (bottom row) light intensity are shown. The traces are zoomed in as for the graphs in Figure 3A. For all three interneurons, at either medium or high light intensity, there was no significant effect of diet on locomotor suppression in the experimental genotype when analyzing either the change in locomotion at light onset (first 1 sec) or during the entire light period (p>0.05, two-way ANOVA with Bonferroni’s post-tests). Flies were one-day starved for all behavioral experiments shown in this figure. Numbers in parentheses after the neuron name denote which split-Gal4 line was used.

Individually activating 5 of the 10 interneuron types caused significant locomotor suppression (Figure 4B and S3B), including 4 of the 7 second-order neurons (Rattle, FMIn, G2N-1, Zorro) and 1 of the 2 third-order neuron types (Fdg). All 5 of these neurons had an effect at multiple light intensities. Some neurons (Rattle, FMIn, Fdg) evoked strong locomotor suppression throughout most of the light stimulation, whereas others (G2N-1, Zorro) had a weaker effect (Figure 4B).

In the positional preference assay, 2 of the 10 interneuron types (Rattle, FMIn) caused significant innate preference (Figure 4C), which was apparent at multiple light intensities (Figure S3C). Both of these neurons were second-order neurons that also affected locomotion. In addition, both types of third-order neurons (Fdg, Bract) caused significant preference during some portion of the preference assay even though the final preference was not significant (Figure 4C and S3C). For Bract, this effect was only observed at a single light intensity for one of the two split-Gal4 lines, suggesting a relatively modest effect. For Fdg, preference was primarily observed with one split-Gal4 line at multiple intensities, although the other line showed a similar trend. Despite the effect of these interneurons on innate preference, none of the 10 interneurons caused significant learned preference (Figure S3D).

Together, these results (Figure 4D) demonstrate both overlap and divergence in the neural circuits that mediate different behavioral responses to sugar. Some second- and third-order neurons can drive several types of behavioral responses, implying that they connect to multiple behavioral pathways, whereas others act specifically to regulate only proboscis extension. The fourth-order neuron, Roundup, did not have a significant effect on any behaviors other than proboscis extension, suggesting that it may have a dedicated role in this behavior. Thus, circuits mediating different innate responses to sugar likely diverge at the level of second- and third-order neurons. Circuits for innate versus learned behaviors may diverge earlier, at the first synapse, given that none of the interneurons elicited associative learning, but we cannot rule out the possibility that we might observe learning effects with different learning protocols or if multiple interneurons were activated simultaneously.

We asked whether the effects of each interneuron on different behaviors were correlated (Figure S3E). The effect of each line on proboscis extension was not correlated with its effect on locomotion, positional preference, or learning at any stimulation intensity. Effects on locomotion and positional preference were correlated for some light intensities, but neither behavior was correlated with learning.

### Connectivity analysis of sugar pathway interneurons

The results described above suggest that second-order sugar neurons are functionally diverse, with some neurons having broader behavioral roles than others. To gain insight into whether this functional diversity of second-order neurons relates to their downstream connectivity, we identified their postsynaptic targets using a recently annotated whole-brain connectome.^42–44^ Individual second-order neurons in one hemisphere generally synapsed onto ∼35-50 postsynaptic neurons with the exception of Usnea, which had over 100 targets. The number of shared targets between different pairs of second-order neurons varied widely, from 0 to 19 neurons (Figure 4E). Phantom and Usnea share many targets with each other but almost no targets with the other second-order neurons. Conversely, Zorro, Rattle, FMIn, and G2N-1 share many targets with each other but very few with Phantom or Usnea. Because Phantom and Usnea are predicted to be inhibitory and the other second-order neurons are predicted to be excitatory,^8^ these analyses reveal a separation between inhibitory and excitatory subcircuits within the sugar pathway. Clavicle is more closely linked to the excitatory subcircuit but shares more targets with Phantom and Usnea than any other excitatory second-order neuron, representing an intermediate role. Analyzing second-to third-order neuron connectivity using output similarity, which takes connection strength into account, revealed the same two subcircuits (Figure S3F) as analyzing the number of shared targets.

The 4 second-order neurons that evoked locomotor suppression (Zorro, Rattle, FMIn, and G2N-1) are the 4 neurons that form the core excitatory subcircuit, suggesting that this subcircuit connects to neurons that regulate locomotion. The neuronal targets shared by these 4 second-order neurons represent candidate neurons mediating locomotor suppression. This list comprises 3 neurons, including Fdg, which also suppressed locomotion. Moreover, Fdg does not receive input from the 3 second-order neurons that failed to significantly affect locomotion (Figure 4F). Thus, Fdg likely represents a critical node in the circuit mediating locomotor suppression to sugar. A recent preprint identified two neurons, Foxglove and Bluebell, that cause locomotor stopping by inhibiting descending neurons that promote walking.^45^ Foxglove and Bluebell both receive input from Fdg, with Fdg’s input to Foxglove being exceptionally strong (105 or 120 synapses depending on the hemisphere). Foxglove and Bluebell also receive direct input from FMIn and Rattle, but not other second-order sugar neurons, and this direct input may explain why FMIn and Rattle elicited much stronger locomotor suppression than Zorro or G2N-1.

Similarly, the neuronal targets shared by Rattle and FMIn, but not other second-order neurons, represent candidate neurons mediating positional preference. Aside from Foxglove and Bluebell, this list comprises 4 neurons, including 2 descending neurons, which have not been functionally characterized. The third-order neuron Bract, which elicited transient preference, receives input from Rattle but not FMIn, nor any of the other second-order neurons (Figure 4F). Bract may contribute to innate preference in parallel with other neurons receiving input from Rattle and/or FMIn.

### Behavioral effects elicited by sugar pathway interneurons are not altered on a high-sugar diet

Having shown that a subset of second- and third-order neurons in the sugar pathway regulate locomotion and innate preference, we asked whether a high-sugar diet blunts their behavioral effects, as we observed with sensory neurons (Figure 3). Observing an effect would suggest that a high-sugar diet affects sugar taste processing in the central brain, in addition to its known effects on sensory neurons.^17, 38, 39^ For the 3 interneurons that strongly affected behavior (Rattle, FMIn, and Fdg), we compared the responses of flies raised on a control versus high-sugar diet. We focused on analyzing locomotion, as this behavior was the most robustly affected by interneuron activation. In contrast to our results with sensory neurons (Figure 3A-B), the locomotor suppression elicited by activating each interneuron was not altered by a high-sugar diet (Figure 4G). These results show that a high-sugar diet acts upstream of second- or third-order neuron activation to alter sugar processing, and thus is likely to act specifically within sensory cells.

## DISCUSSION

Using sugar taste as a model, we investigated how neural circuits are organized to elicit multiple behavioral responses to a single sensory stimulus. We first asked how different behavioral responses to sugar are modulated by internal and external cues. In general, different behaviors were coordinately regulated, showing enhancement by hunger and suppression by bitter taste or a high-sugar diet, but the degree of modulation varied. We then asked whether neurons known to mediate the initiation of feeding also regulate other behavioral responses to sugar. We found that circuits mediating different innate responses to sugar partially overlap and diverge at the level of second- and third-order neurons, whereas circuits for innate versus learned behaviors may diverge at the first synapse. Connectomic analyses revealed distinct subcircuits for sugar processing within the SEZ, providing a potential anatomical substrate for our behavioral results.

### Behavioral responses to sugar neuron activation

Optogenetic activation of sugar neurons suppressed locomotion and elicited innate and learned preference, reproducing known effects of sugar.^18–21^ However, one counterintuitive result was that innate preference diminished and switched to avoidance with higher intensity stimulation, an observation also noted in a previous study.^26^ With natural sugar, positional preference increases with concentration even up to very high concentrations such as 2M sucrose.^19^ One possibility is that optogenetically induced avoidance results from unnatural levels or patterns of sugar neuron activation, which may cause compensatory changes in the downstream circuit.

Alternatively, different subsets of *Gr64f-Gal4*-expressing neurons may promote opposing behaviors, as suggested by a recent study,^46^ and high levels of activation may preferentially recruit aversive circuits. While it is possible that aversive *Gr64f-Gal4*-expressing neurons represent off-target cells that do not endogenously express Gr64f, flies naturally avoid sugar in some contexts, such as egg-laying, and subsets of sugar-sensing neurons have been shown to elicit aversion.^47, 48^ Neither of the downstream neurons that elicited positional preference (Rattle and FMIn) elicited aversion at high light intensity, suggesting that high intensity stimulation of *Gr64f-Gal4*-expressing neurons elicits aversion through a distinct pathway.

All of the behavioral effects of sugar neuron activation were strongly modulated by hunger state, contrasting with the minimal effect of hunger on bitter-evoked behaviors. This differential modulation aligns with the ethological relevance of each taste modality. Sweet taste promotes caloric intake, which is only necessary when energy stores are low. Bitter taste prevents the ingestion of toxins, which is important whether or not an animal is food-deprived. When we simultaneously activated sugar- and bitter-sensing neurons, the effects of bitter neurons dominated over the effects of sugar neurons. The taste circuit likely evolved this strict hierarchy due to the potentially deadly consequences of ingesting toxins; preventing such an outcome should generally take precedence over caloric consumption. How this gating of sugar input by bitter input occurs at the circuit level will be an interesting question for future study.

Although hunger strongly influenced all sugar-evoked behaviors, different behaviors were not equally affected. Hunger also acted selectively in modulating bitter-evoked behaviors: it enhanced learned aversion but did not affect locomotion or innate preference. Moreover, previous studies have shown that hunger suppresses the inhibitory effect of bitter on proboscis extension,^13, 29^ which may encourage energy-depleted flies to consume contaminated food sources in order to avoid starving. These results argue that behaviors elicited by the same taste can be differentially gated by hunger, even in opposing directions. To achieve this selective modulation, hunger-dependent gating must occur downstream of sensory neurons. Some previous studies observed hunger modulation of sugar and bitter sensory neuron responses,^13, 24, 29^ but additional modulation likely occurs in the downstream circuit.^8, 49, 50^

### Effects of a high-sugar diet on sweet taste processing

A high-sugar diet reduces proboscis extension to sugar but leads to increased long-term feeding.^15, 17, 39^ We found that a high-sugar diet reduces other behavioral responses to sugar, suggesting that short-term responses to sugar are globally dampened. How and where does this suppression occur? Previous studies found that sugar-enriched diets reduce sugar-evoked activity and synaptic output of sugar sensory neurons.^17, 33, 51^ Reduced activity has been observed in recordings of taste sensilla,^17, 33^ where the dendrites and cell bodies of sensory neurons reside, suggesting that high-sugar diets may desensitize or downregulate sugar receptors or reduce neuronal excitability, thus limiting the depolarization elicited by sugar receptor currents. Transcriptomic studies found that high-sugar diets did not significantly alter the expression of sugar receptors,^33, 38^ but these results do not rule out changes in receptor sensitivity or function. Because our experiments used optogenetic stimulation of sugar neurons, bypassing sugar receptor activation, our results conclusively demonstrate that a high-sugar diet modulates sugar responses independently of any effect on sugar receptors.

We expect that changes in the excitability and firing of sugar-sensing neurons contribute to the effects of a high-sugar diet, but we wondered whether diet may also alter neural processing in the downstream circuit. Indeed, excess dietary sugar leads to extensive transcriptional changes in the brain, many of which are distinct from transcriptional changes in taste organs.^33^ We tested whether a high-sugar diet modulated behavioral responses elicited by activating second- or third-order sugar neurons, thus bypassing the sensory neurons. The high-sugar diet did not reduce these behavioral responses, suggesting that it alters sugar processing by acting specifically within sensory neurons.

### Divergence of neural circuits for different behaviors

Where and how does a sensory circuit diverge into different pathways that mediate different behaviors? If the sensory stimulus is complex and requires extensive processing to be identified, this divergence may occur after several layers of sensory processing. This may be why extensive processing of visual input occurs in the periphery before visual pathways for different functions begin to diverge in the brain.^52^ In contrast, a clear divergence emerges very early in the olfactory system: at the level of second-order neurons, which project to multiple brain regions with distinct behavioral roles.^53^ Consequently, most olfactory processing occurs within each downstream region and presumably fulfills specific functions related to each area’s role. Taste is often considered to be a relatively simple stimulus, suggesting that pathways for different taste-evoked behaviors may diverge early in the circuit. In mammals, surprisingly little is known about how taste-sensing pathways interface with behavioral circuits. The canonical taste pathway relays taste information from the tongue to the gustatory cortex via the brainstem and thalamus,^54^ and different behavioral pathways may diverge within the cortex. For example, projections from the gustatory cortex to the amygdala regulate innate licking behavior but are dispensable for associating taste identity with a learned action.^55^ Projections from the gustatory cortex to the orbitofrontal cortex may be involved in more complex feeding decisions.^54, 56^

Studies in *Drosophila*, where the architecture and function of neural circuits can be mapped at the level of single cells, are well-suited to provide insight into how and where neural circuits diverge into different pathways that regulate distinct behaviors. Our experiments reveal that circuits mediating different innate responses to sugar – including proboscis extension, locomotor suppression, and positional preference – involve partially overlapping sets of second- and third-order neurons but distinct fourth-order neurons. We propose that basic aspects of taste processing occur within second-order neurons, such as pooling information across sensory cells, while processing in third-order neurons begins to reflect sensorimotor transformations required for specific behaviors.

Our results also suggest that circuits for innate versus learned taste behaviors may diverge at the first synapse, although we cannot rule out the possibility that different learning protocols or co-activation of multiple interneurons might reveal overlapping circuits for innate and learned responses. Associative learning requires the mushroom body,^7^ located in the superior protocerebrum, so sugar input used for learning must be transmitted out of the SEZ to the mushroom body. All of the interneurons in the proboscis extension circuit reside solely within the SEZ,^8^ but other second-order sugar neurons project to the superior protocerebrum.^57^ Kim et al. (2017) identified a second-order neuron that conveys sugar input to the mushroom body and is required for flies to learn sugar-bitter associations that modulate proboscis extension to sugar.^58^ It is unclear whether this neuron is also involved in odor-sugar associations, where sugar serves as the unconditioned stimulus rather than the conditioned stimulus. In the future, further connectomic and functional studies of downstream taste circuits should reveal further insight into the circuits that mediate distinct behavioral responses to sugar.

## AUTHOR CONTRIBUTIONS

A.V.D conceived and supervised the project, conducted proboscis extension experiments, and performed connectome analyses. R.V.J, C.X.W., F.V.L.-P., L.N., H.A.U., and J.U.D. conducted all other behavioral experiments. R.V.J., F.V.L.-P., and C.X.W. performed brain dissections and immunostaining to confirm Gal4 expression patterns. A.V.D. generated figures and wrote the manuscript with input from all authors.

## ACKNOWLEDGMENTS

We thank Richard Axel for his support during initial stages of this project, Jan Hawes for facilities assistance, Maia Yang for assisting with confirmation of Gal4 expression patterns, and Gabriella Sterne and Chris Rodgers for comments on the manuscript. We are grateful to Gabriella Sterne, the Janelia Research Center, and the Bloomington Drosophila Stock Center (BDSC) for providing fly strains. We acknowledge the Princeton FlyWire team and members of the Murthy and Seung labs for development and maintenance of FlyWire (supported by BRAIN Initiative grant MH117815 to Murthy and Seung). This work was supported by the Whitehall Foundation (A.V.D.).

## DECLARATION OF INTERESTS

The authors declare no competing interests.

## STAR METHODS

## RESOURCE AVAILABILITY

### Lead contact

Further information and requests for resources and reagents should be directed to and will be fulfilled by the lead contact, Anita Devineni.

### Materials availability

This study did not generate new unique reagents.

### Data and code availability

- All data reported in this paper will be shared upon request to the lead contact.
- This study did not generate original code.
- Any additional information required to reanalyze the data reported in this paper is available from the lead contact upon request.

## EXPERIMENTAL MODEL AND SUBJECT DETAILS

### Fly strains and maintenance

Flies were reared at 25°C on standard cornmeal or cornmeal/molasses food and maintained in constant darkness to avoid activation of Chrimson. All experiments used mated females, which were collected within a few days after eclosion. Flies were placed on food containing 1 mM all trans-retinal for 3-4 days prior to testing. During starvation, flies were housed in vials with 1 mM all trans-retinal solution on a piece of Kimwipe. The high-sugar diet consisted of cornmeal/molasses food with 30% sucrose (wt/vol) added.

Genotypes used for each experiment are specified in the figures and legends, and detailed information regarding fly strains is provided in the Key Resources Table. For interneuron activation, multiple split-Gal4 lines were available for some lines. We selected initial lines for testing based on the strength of their proboscis extension phenotype in Shiu, Sterne et al.^8^ and the specificity of their expression pattern, based on images in the Janelia FlyLight Split-Gal4 database (https://splitgal4.janelia.org/) acquired by Sterne et al.^41^ For neurons that elicited borderline phenotypes in any behavioral assay, we tested a second line labeling the same neuron (if available).

## METHOD DETAILS

### Behavioral assays

Assays for locomotion, innate preference, and associative learning were conducted using previously described methods.^28^ The behavioral arena contains a circular chamber with a glass cover, and flies are filmed from above using a USB camera. The infrared light for illumination and the red (627 nm) LED array for optogenetic stimulation are located just beneath the chamber. 20-25 flies were tested per trial. Flies were loaded into the chamber using mouth aspiration and were given ∼3 minutes to habituate before the experiment started. Flies were filmed at 30 frames/second.

Locomotion and innate preference were assayed sequentially in the same flies. To quantify locomotor effects, light stimulation was presented for 5 sec. Each of the three light intensities was delivered to the same flies in increasing order of intensity, with several minutes between stimulations to ensure that the behavior had recovered. After an additional rest period, the same flies were then tested for innate preference by delivering low intensity light stimulation to two opposing quadrants for 30 sec. The flies then had a 30 sec rest period without light, followed by low intensity light stimulation in the other two quadrants for 30 sec. Switching the light quadrants ensured that we could assess light preference independently of any spatial bias. This protocol was then repeated sequentially with medium and high intensity light.

For associative learning, the odors used were 3-octanol (1:1000) and 4-methylcyclohexanol (1:750). The flow rate was 100 mL/min. First, the CS+ odor was presented in all quadrants for 1 min along with 1 Hz light stimulation that started 5 sec after odor valve opening, following the protocol used in previous learning studies.^27^ After a 1 min rest, the CS-odor was presented alone for 1 min. Following another 1 min rest, the CS+ and CS-odors were simultaneously delivered to different sets of opposing quadrants for 1 min, allowing the flies to choose between the odors. After another 1 min rest, the two odors were presented again for 1 min but the odor quadrants were switched to control for any spatial bias. Which odor was used as the CS+ or CS-was counterbalanced across trials. The light intensity used for learning experiments with *Gr64f-Gal4* was 4 µW/mm^2^ (the intensity that elicited preference in the innate choice assay), but we also tested 30 µW/mm^2^ in Figure S1B. 30 µW/mm^2^ was used to activate *Gr5a*-expressing neurons (Figure S1G) or downstream sugar circuit interneurons (Figure S3D).

Fly videos were analyzed using FlyTracker,^59^ and FlyTracker output was further analyzed in MATLAB to quantify locomotion or preference. Preference index (PI) values were quantified in 1 sec bins. For innate preference, PI was calculated as (# flies in light quadrants – # flies in non-light quadrants) / total # flies. For learning, PI for the conditioned odor was calculated as (# flies in CS+ quadrants – # flies in CS-quadrants) / total # flies. Forward and angular velocities were averaged over 0.33 sec bins (10 frames). To quantify locomotor changes at light onset or throughout the light period, we averaged forward or angular velocity over the first 1 sec or entire 5 sec of light presentation, respectively, and subtracted these values from the baseline values averaged over the 4 sec preceding light onset. For statistical analyses of these data, each trial was considered to be a single data point (“n”).

Proboscis extension assays were conducted in one-day starved flies as previously described.^49, 60^ Flies were immobilized on their backs with myristic acid, and the two anterior pairs of legs were glued down so that the proboscis was accessible. Flies recovered from gluing for 30-60 minutes in a humidified chamber and were water-satiated before testing. To activate taste neurons with Chrimson, a red LED (617 nm) was manually turned on for 1 second. Each fly was given two chances to respond to a light stimulus. At the end of each experiment, flies were tested with a positive control (500 mM sucrose) and were excluded from analysis if they did not respond.

### Connectome analysis

Annotated neuronal connectivity from a whole-brain dataset^42–44^ was downloaded from FlyWire using the latest release (Snapshot 630). Each of the sugar circuit interneurons has been previously identified and labeled in this connectome.^8^ To examine the targets and output similarity between second-order neurons, and to make the diagram in Figure 4F, we focused on neurons in a single hemisphere (left). To determine which second-order neurons provide input to Fdg, Bract, Foxglove, or Bluebell, we analyzed inputs to cells in both hemispheres. Cell IDs are: Zorro (720575940629888530), Rattle (720575940638103349), FMIn (720575940614763666), G2N-1 (720575940620874757), Clavicle (720575940655014049), Phantom (720575940616103218), Usnea (720575940632648612), Fdg (720575940631997032, 720575940647030324), Bract (720575940645045527, 720575940610001220, 720575940626557442, 720575940627285267), Foxglove (720575940611160930, 720575940628755815) and Bluebell (720575940631168057, 720575940635166571). To quantify the output similarity between two neurons we used cosine similarity, a metric that has previously used in connectomic analyses.^61^ We calculated the cosine of the angle between the vectors containing the number of synapses onto each output neuron. These vectors are n-dimensional, where n is the total number of output neurons for the two input neurons.

## QUANTIFICATION AND STATISTICAL ANALYSIS

Statistical analyses were performed using GraphPad Prism, Version 9. Statistical tests and results are reported in the figures and legends. Most experiments had three genotypes, so we used one-way ANOVA followed by Dunnett’s test comparing experimental flies to each control. In cases where only two groups were being compared, we used unpaired t-tests. To test the effect of sugar neuron activation on different diets, we used two-way ANOVA followed by Dunnett’s test comparing experimental flies to genetic controls or Bonferroni’s test comparing flies on each diet. For proboscis extension experiments, which generate binary data, we used Fisher’s exact test to compare experimental and control flies. All graphs represent mean ± SEM. Sample sizes are listed in the figure legends.

## KEY RESOURCES TABLE

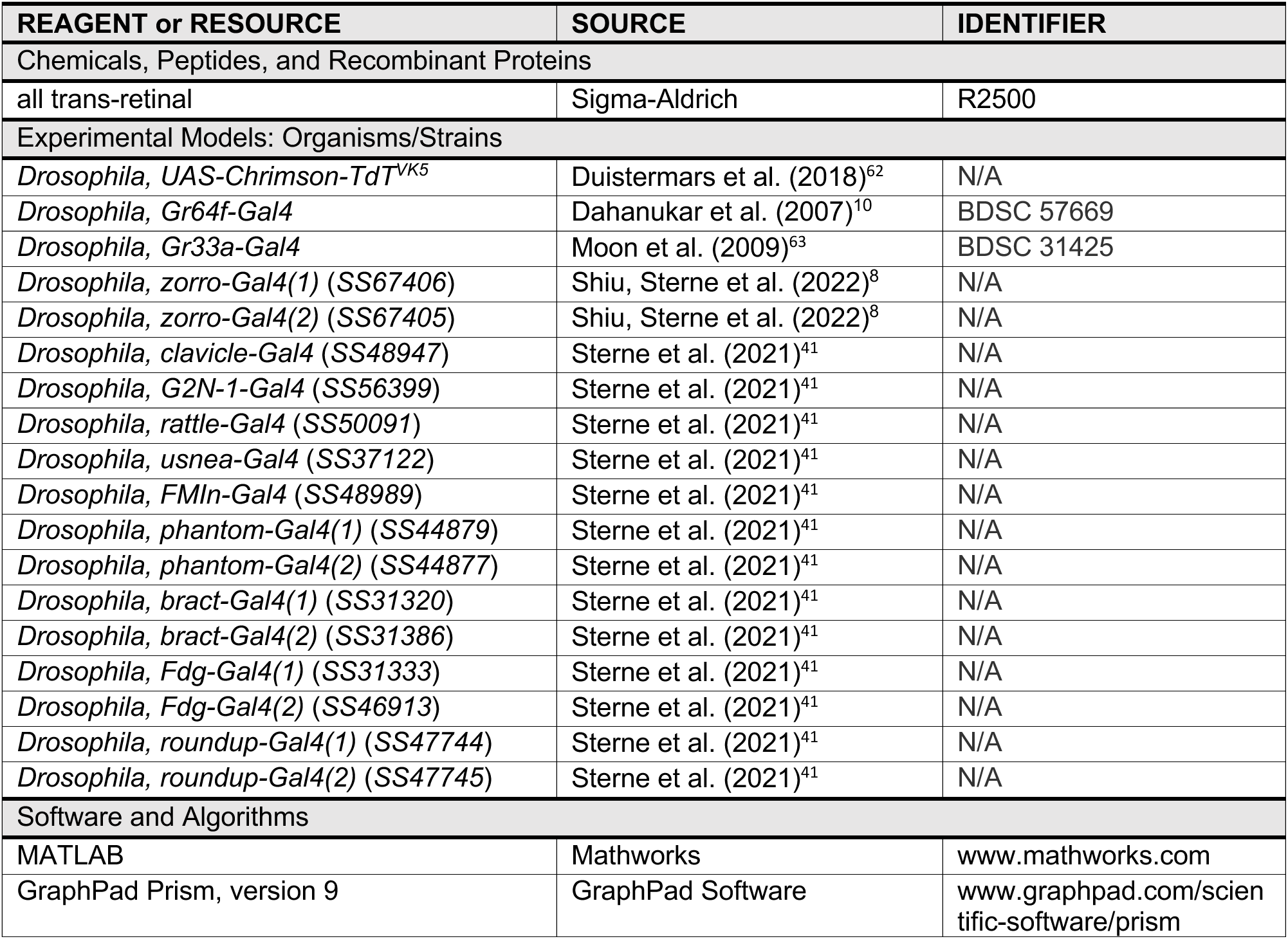

---

**Figure S1, related to Figure 1.**
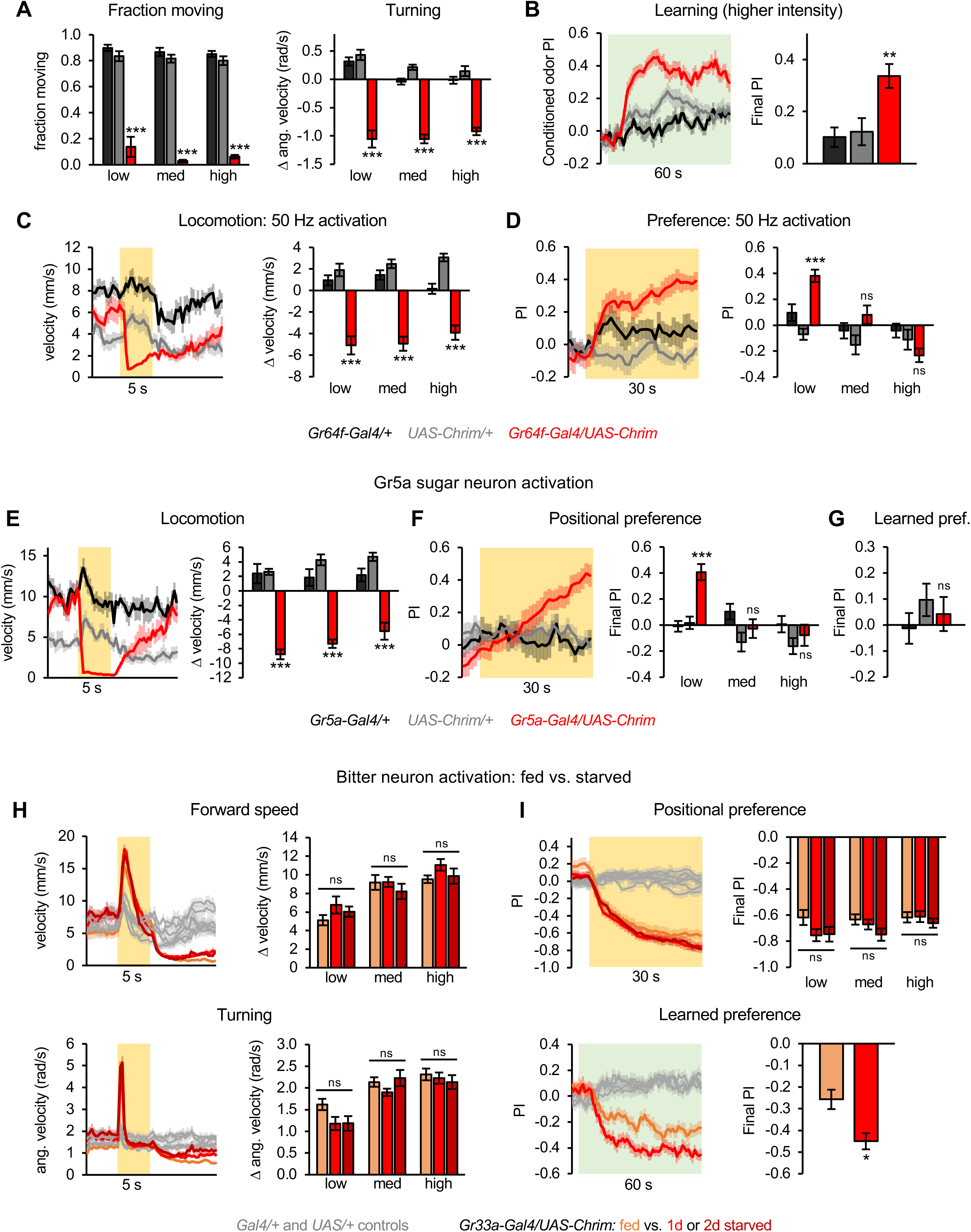
Additional characterization of effects induced by sugar activation and comparison to bitter activation. (A) Sugar neuron activation in fed flies decreased the fraction of flies moving (left) and angular velocity (right) (n = 12 trials). Values are quantified for the first 1 sec of light stimulation. (B) Activating sugar neurons at a higher light intensity than in Figure 1 (30 µW/mm^2^) elicited learned odor preference (n = 24 trials, 12 sets of flies). (C-D) Activating sugar neurons using 50 Hz pulsed light in starved flies elicited similar effects on locomotion (C; n = 8-9 trials) and positional preference (D; n = 12-18 trials, 6-9 sets of flies) as those observed with continuous light (Figure 1). (E-G) Activation of sugar-sensing neurons using *Gr5a-Gal4* elicited locomotor suppression (E; n = 8 trials) and positional preference (F; n = 16 trials, 8 sets of flies) but not learned preference (G; n = 24 trials, 12 sets of flies). (H-I) Activation of bitter-sensing neurons using *Gr33a-Gal4* elicited aversive behaviors that are similar in fed and starved flies, with the exception of stronger aversive learning in starved flies. Data for fed flies are the same data shown in Deere et al., 2023. (H) Locomotor effects elicited by bitter neuron activation in fed and starved flies. Traces show forward and angular velocity for medium intensity light stimulation (n = 10-12 trials). Bar graphs quantify the change in these parameters at light onset. (I) Innate positional aversion (top; n = 20-24 trials, 10-12 sets of flies) and learned aversion (bottom; n = 22-24 trials, 11-12 sets of flies) elicited by bitter neuron activation in fed and starved flies.

**Figure S2, related to Figure 3.**
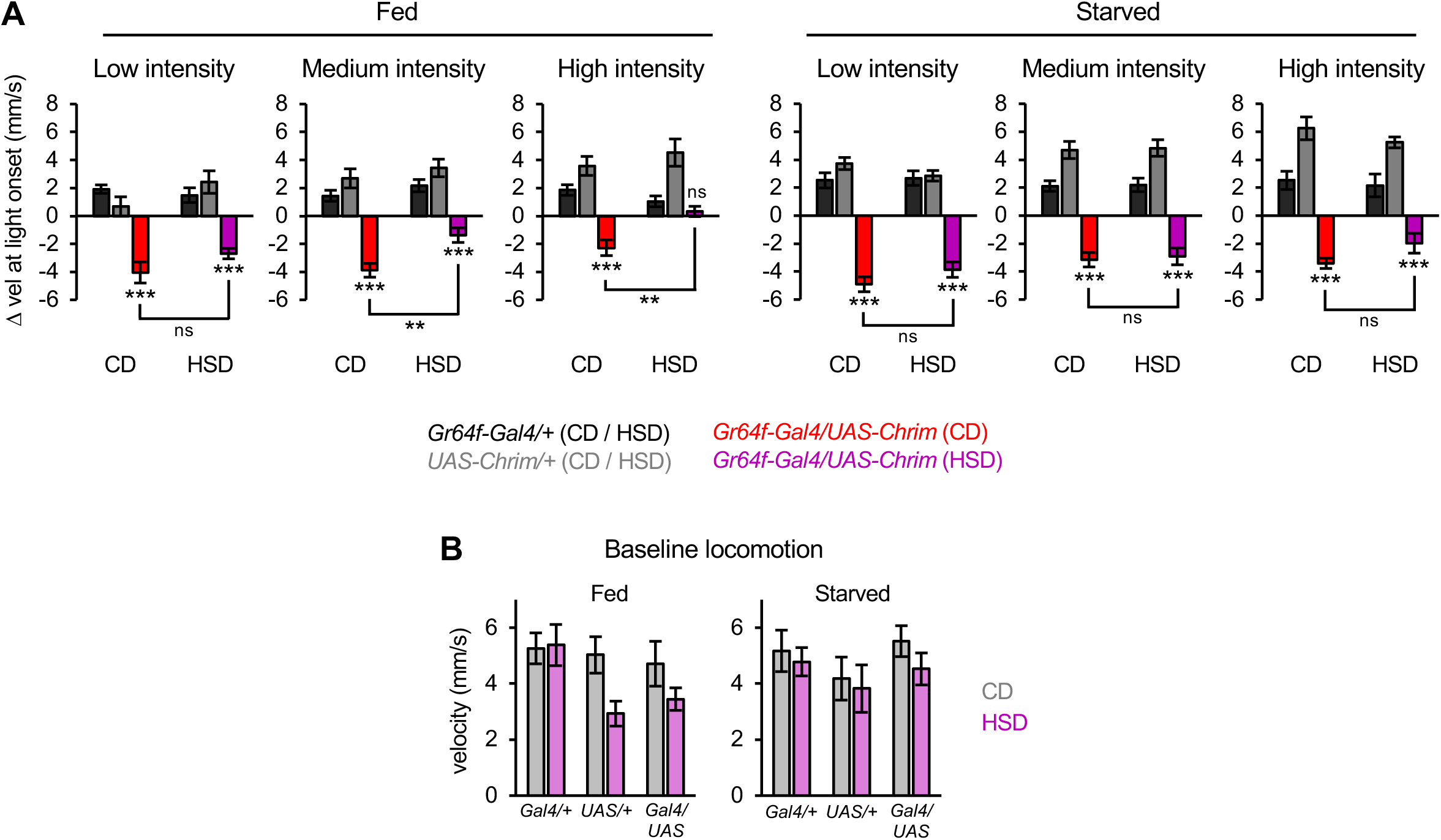
Additional characterization of behavioral responses on a high-sugar diet. (A) Change in velocity at light onset (first 1 sec) quantified from the data in Figure 3A (n = 7-10 trials). (B) Baseline locomotion for sugar activation experiments shown in Figure 3A (n = 7-10 trials). In fed flies, there was a significant overall effect of diet (p<0.05) but no significant effect for any individual genotype. There was no significant effect of diet in starved flies (p>0.05). All statistics in this figure represent the results of two-way ANOVA followed by Dunnett’s post-tests comparing experimental flies to each control or Bonferroni’s post-tests comparing CD and HSD.

**Figure S3, related to Figure 4.**
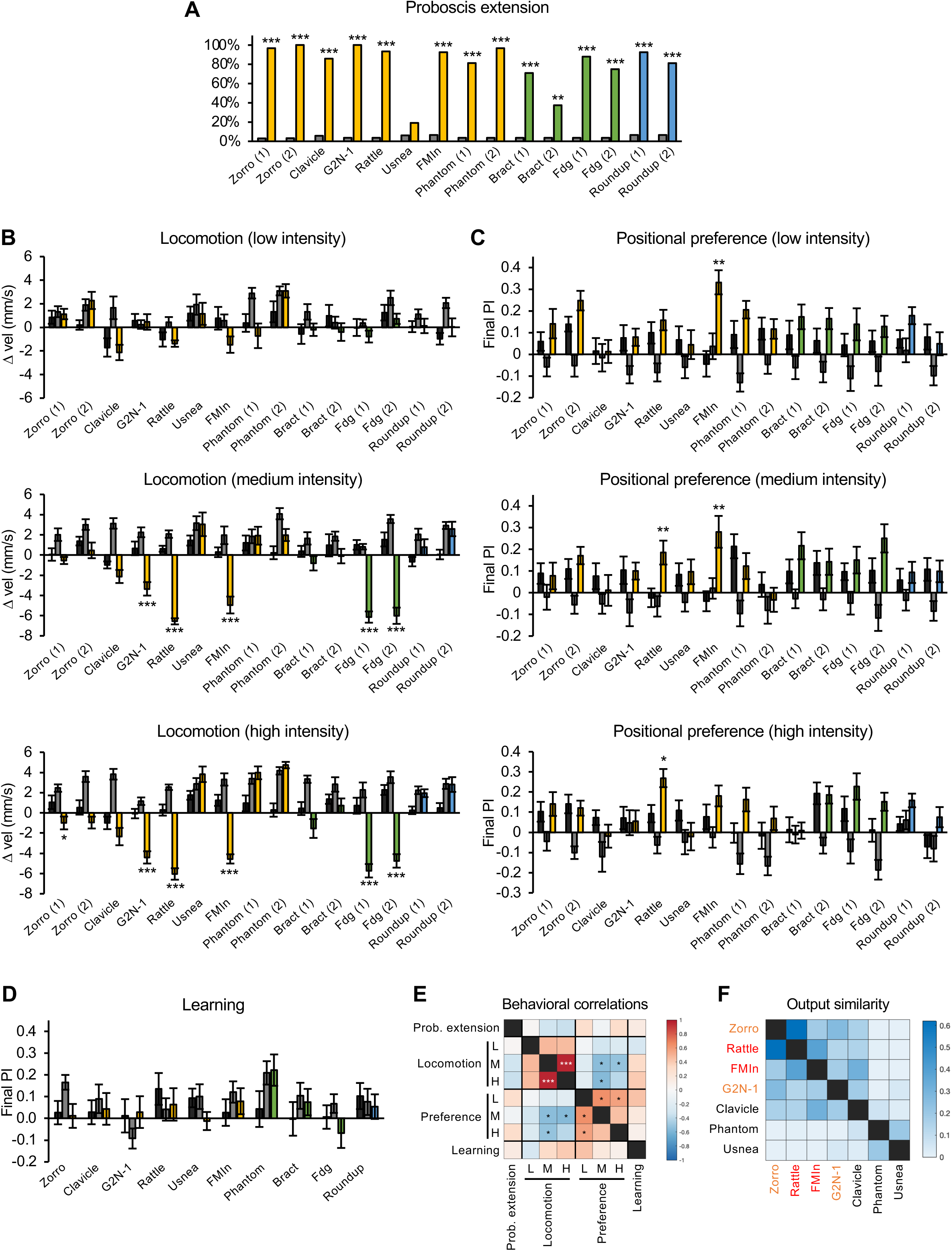
Additional characterization of behavioral effects elicited by sugar circuit interneurons. (A) Proboscis extension elicited by activating each type of interneuron in the sugar circuit (Fisher’s exact test, n = 25-47 flies). (B) Change in forward speed (first 1 sec) elicited by activation of each type of sugar circuit interneuron at each light intensity (n = 7-12 trials). (C) Innate positional preference elicited by activation of each type of sugar circuit interneuron at each light intensity (n = 14-24 trials, 7-12 sets of flies). (D) Learned odor preference elicited by activation of each type of sugar circuit interneuron (n = 16-24 trials, 8-12 sets of flies). (E) Correlation values (Pearson’s r) between different behavioral effects elicited by activating sugar pathway interneurons. Data from all lines tested in Figure S3 are included in the analysis (n=10 for correlations with learning and n=15 for all other correlations). Values for “locomotion” represent the change in forward speed during the first 1 sec of light, so negative values represent locomotor suppression. (F) Heatmap showing the cosine similarity of outputs for each pair of second-order neurons. Cosine similarity ranges from 0 to 1. Neurons are colored red if they affected both locomotion and preference and orange if they only affected locomotion. In panels A-D, flies were one-day starved. Black and grey bars represent the Gal4/+ and UAS/+ controls, respectively, and colored bars represent experimental flies carrying the split-Gal4 and UAS-Chrim. Second-order neurons are listed in descending order of the number of synapses they receive from sugar neurons (as reported in Shiu, Sterne et al., 2022). Numbers in parentheses after the neuron name denote which split-Gal4 line was used.

## Notes

### Competing Interest Statement

The authors have declared no competing interest.

